# Are you talking to me? How the choice of speech register impacts listeners’ hierarchical encoding of speech

**DOI:** 10.1101/2024.09.02.610805

**Authors:** Giorgio Piazza, Sara Carta, Emily Ip, Jose Pérez-Navarro, Marina Kalashnikova, Clara D. Martin, Giovanni M. Di Liberto

## Abstract

Speakers accommodate their speech to meet the needs of their listeners, producing different speech registers. One such register is Foreigner-Directed Speech (FDS), which is the way native speakers address non-native listeners, typically characterized by features such as slow speech rate and phonetic exaggeration. Here, we investigated how register impacts the cortical encoding of speech at different levels of language integration. Specifically, we tested the hypothesis that enhanced comprehension of FDS compared with Native-Directed Speech (NDS) involves more than just a slower speech rate, influencing speech processing from acoustic to semantic levels. Electroencephalography (EEG) signals were recorded from Spanish native listeners, who were learning English (L2 learners), and English native listeners (L1 listeners) as they were presented with audio-stories. Speech was presented in English in three different speech registers: FDS, NDS and a control register (Slow-NDS) which is slowed down version of NDS. We measured the cortical tracking of acoustic, phonological, and semantic information with a multivariate temporal response function analysis (TRF) on the EEG signals. We found that FDS promoted L2 learners’ cortical encoding at all the levels of speech and language processing considered. First, FDS led to a more pronounced encoding of the speech envelope. Second, phonological encoding was more refined when listening to FDS, with phoneme perception getting closer to that of L1 listeners. Finally, FDS also enhanced the TRF- N400, a neural signature of lexical expectations. Conversely FDS impacted acoustic but not linguistic speech encoding in L1 listeners. Taken together, these results support our hypothesis that FDS accommodates speech processing in L2 listeners beyond what can be achieved by simply speaking slowly, impacting the cortical encoding of sound and language at different abstraction levels. In turn, this study provides objective metrics that are sensitive to the impact of register on the hierarchical encoding of speech, which could be extended to other registers and cohorts.

## 1. Introduction

When addressing second language (L2) learners, L1 speakers naturally tend to speak in a particularly clear manner by using a speech register known as Foreigner Directed Speech (FDS; Scarborough et al., 2007; Tarone, 1980; Uther et al., 2007; see Piazza et al., 2022 for a review of the acoustic features of FDS). FDS is often studied in comparison with Native Directed Speech (NDS), which is the register used between L1 speakers, without the intention of enhancing intelligibility (Ferguson & Kewley-Port, 2002). FDS has also been referred to as “non- native directed speech” (NNDS; e.g., Piazza et al., under review; 2023) since it is not limited to foreign listeners, and as “L2 speech accommodation” because it is assumed to be the result of the speaker’s accommodation to the listener’s low L2 proficiency and learning needs (Giles, 2016; Lindblom, 1990; Zhang & Giles, 2017). Few studies investigated directly the impact of FDS use on L2 perception, comprehension, and learning (Piazza et al., 2022, 2023; Uther et al., 2012; see Rothermich et al., 2019 for a review on impressions about FDS). For example, Piazza et al. (2023) provided evidence of the positive impact of FDS on L2 perception and production in a novel L2 word learning task. However, controlled manipulations such as novel word learning do not reflect listeners’ naturalistic exposure to L2 speech, and it remains unknown whether FDS supports perception and comprehension of continuous speech. Also, most studies that investigated FDS perception (with controlled manipulations) employed behavioural experiments (Piazza et al., 2023; Bobb et al., 2019; Kangatharan et al., 2023; but see Uther et al., 2012), which can only provide indirect measures of L2 processing when it is already concluded. Instead, using neurophysiological techniques (e.g., electroencephalography (EEG)), enables the exploration of speech perception as it unfolds. Here, we probed the encoding of a hierarchy of linguistic information in speech and language with EEG to explore whether and how FDS promotes L2 learners’ processing of L2 sounds and discourse.

Accurate multidimensional models of L2 perception require understanding how speech registers and acoustic features modulate brain mechanisms underlying L2 acquisition, including speech perception and comprehension in naturalistic listening task. One way to inform such models with ecological validity is by using EEG to measure cortical encoding (CE) of speech features. We interpret CE as encompassing a broad range of speech features, both continuous and non-continuous, and includes cortical tracking of the speech envelope—a dynamic alignment of brain activity with the temporal modulations of speech (Giraud & Poeppel, 2012; Kalashnikova et al., 2018; Obleser & Kayser, 2019). Cortical tracking of speech is widely regarded as a marker of the linguistic processes involved in speech perception (Giraud & Poeppel, 2012; Luo & Poeppel, 2007; Meyer, 2018), which is linked to enhanced speech clarity and comprehension, particularly in L1 adult listeners (Ahissar et al., 2001; Ding & Simon, 2014; Etard & Reichenbach, 2019; Keitel et al., 2018; Molinaro & Lizarazu, 2018; Peelle et al., 2013; Riecke et al., 2018; but see Peña & Melloni, 2012 for opposite results).

The speech envelope is the low-frequency amplitude modulation of the broadband speech signal, which carries acoustic information important for perceptual and linguistic encoding (Attaheri et al., 2022). Speech envelope encoding is a broad measure of speech perception, which is influenced by factors such as attention and engagement (Ding & Simon, 2014; Keitel et al., 2018; O’Sullivan et al., 2015). Beyond the speech envelope, a growing number of studies are investigating CE of speech at higher order phonological and semantic levels (Brodbeck et al., 2022; Broderick et al., 2018, 2021; Pérez-Navarro et al., 2024). It is possible to do so by mapping specific sets of speech features, both acoustic and abstract, to brain signals (Crosse et al., 2016; Di Liberto et al., 2015, 2018, 2021). Some features of continuous speech have been studied in the EEG response signals, usually within low-frequency bands (<8Hz), by using encoding model estimations derived, for example, with multiple regression. Here we adopt the multivariate Temporal Response Function approach (TRF; Crosse et al., 2016; Di Liberto et al., 2015), which has been shown to enable the study of speech perception with tasks involving continuous speech listening, probing the CE of both speech sounds and linguistic properties, such as the CE of the acoustic envelope (Kalashnikova et al., 2018), phonological properties (Brodbeck et al., 2018; Di Liberto et al., 2015; 2021; 2023; Gillis et al., 2023) and semantic expectation (Broderick et al., 2018, 2021; Klimovich-Gray et al., 2023). Previous studies have shown that it is possible to investigate phonemic processing and categorization across various participant cohorts (Carta, et al., under review; Di Liberto, et al., 2015; 2018; 2021; Klimovich-Gray et al., 2023). Within continuous speech, responses to phonetic features of speech and phonemic categorical processing can be discriminated in the low-frequency EEG signals (Di Liberto et al., 2015). This shows that it is possible to employ TRF to assess L2 learners’ phonological perception, which could be extended to the assessment of how such phonological perception changes depending on the speech register (FDS or NDS). Using a similar approach allowed us to measure phonological processing across different speech registers such as FDS and NDS.

Previous CE research involves the use of semantic prediction features built with computational models estimating how semantically surprising words are given their preceding context (e.g., large language models). For instance, Klimovich-Gray et al. (2023) built a multivariate model that accounted for both speech envelope and semantic surprisal. The TRF weights results of the semantic surprisal regressor highlighted a TRF complex, with prominent centro-parietal negativity, comparable to the classic semantic N400 (Broderick et al., 2018, 2022). For the present study’s purposes, we combine comprehension questionnaire and EEG-based semantic CE measures to test whether FDS promotes comprehension in L2 learners. Furthermore, we carry out multivariate analyses to disentangle the cortical processing of speech and language at the level of acoustics, phonology, and semantics, investigating whether and how that processing is modulated by speech registers.

In addition, previous studies investigated CE of speech registers such as infant directed speech (IDS), which is used to address infants and support their language acquisition (Burnham et al., 2015; Kalashnikova et al., 2017; Kalashnikova & Burnham, 2018; Kuhl, 1997; Trainor & Desjardins, 2002) and share some acoustic features with FDS (Piazza et al., 2022). Research on IDS observed that infants – not adults – exhibited better CE of IDS as compared to other speech registers (Kalashnikova et al., 2018; Menn et al., 2022; see also Attaheri et al., 2022 for similar research). While acknowledging the inherent distinctions between adults and infants, these findings indicate that listeners’ CE is enhanced when listeners are exposed to speech registers specifically intended for them. These results, along with acoustic features and didactic function analogies drawn between IDS and FDS, suggest that L2 learners may benefit from being exposed to FDS. However, these studies typically investigated CE of speech envelope and prosody contours, which represent only one (although important) aspect of speech processing that serves as a proxy of speech encoding. So far, there is a lack of studies that directly measured cortical encoding across speech registers and encoding of various language features (both acoustic and abstract).

Particularly relevant to our study, Verschueren and colleagues (2022) investigated (native) linguistic speech processing as a function of varying speech rate. Their findings showed that slower speech rate led to an increase in CE, suggesting a connection between how linguistic representations are tracked and the reduction in speech rate. Given that the differences in perception between FDS and NDS might arise from differences in speech rates (FDS being slower than NDS), and that speech rate affects CE, in this study we decided to also investigate CE of an artificial speech register serving as a control condition, which we call Slow-NDS. This speech register had the same acoustic features of NDS but a speech rate that was made similar to FDS (slower than NDS) by means of dynamic time warping (Müller, 2007). We expected enhanced speech perception in L2 learners to be mainly due to acoustic feature accommodation of FDS (Bobb et al., 2019; Piazza et al., 2022, 2023), as opposed to speech rate.

Here, we investigated how the exposure to speech register impacts the CE of speech across several levels of the processing hierarchy, by probing acoustic, phonological, and semantic processing with EEG and multivariate encoding models. We presented two EEG experiments involving English L2 learners (Spanish L1, henceforth L2L) and English native speakers (English L1, henceforth L1L). During these experiments, EEG signals were acquired as participants listened to continuous speech (stories), and were asked comprehension questions. Stories were presented in three different speech registers: FDS, NDS, and Slow- NDS. We expected CE to be enhanced in L2 learners exposed to FDS relative to both NDS and Slow-NDS at various levels of the hierarchy (from acoustics to semantics). We also expected a facilitatory effect of slow speech rate, thus CE to be more enhanced in Slow-NDS than NDS. Conversely, since L1 listeners are not the intended addressees of FDS, which is not accommodated to promote L1 speech perception, we hypothesized that slow speech rate in FDS and Slow-NDS would only enhance CE of speech envelope, as compared to NDS. In addition, we predicted FDS to promote L2L’s phonological perception as compared to both Slow-NDS and NDS. Then, for L2L we expected both higher comprehension scores and increased semantic CE for FDS as compared to Slow-NDS and for Slow-NDS as compared to NDS. Conversely, L1L, who have native proficiency of English, were expected to show close-to-ceiling performance in understanding all stories. That is, they were not expected to benefit from the exposure to any speech register in their comprehension accuracy and encoding of semantic surprisal. Here, we aimed to elucidate whether high level metrics of cortical encoding can be used to assess CE differences across speech registers.

## 1. Method

### 2.1. Participants

#### 2.1.1. Experiment 1 (L2L)

A total of 28 participants, aged between 18-35, were recruited to take part in experiment 1. They were L2 learners of English (L1 speakers of Spanish), with mid-low proficiency in English (henceforth L2L; M_age_ = 22.8 y.o., SD = 3.42, Female = 21). L2L participants were tested for their English level in an individual interview with an expert linguist, who assigned marks from 0.0 to 5.0 (0.0 = no knowledge; 1.0 = low; 5.0 = native-like). In the interview, fluency, vocabulary, grammar, and pronunciation were evaluated, and altogether concurred in the overall mark. We only recruited participants who obtained an overall mark between 1.0 and 3.0 (M = 2.96, SD = 0.34). Of the original L2L sample, 2 participants were excluded due to technical problems and 1 due to very low comprehension score (4% of correct responses), leaving the final cohort to 25 L2L. The experiment was carried out at the Basque Center on Cognition, Brain and Language (Spain). The study was approved by the BCBL Ethics Committee. All participants signed an informed consent form prior to the experiment. Participants were paid 20 euro for taking part in the study.

#### 2.1.2. Experiment 2 (L1L)

Twenty-seven native speakers of English (L1L), aged between 18-31, were recruited to take part in the Experiment 2 and were tested at Trinity College Dublin (Ireland) (M_age_ = 22.15, y.o., SD = 3.05, Female = 13). Two participants were excluded due to a technical issues, leaving the final cohort to 25 participants. Of these, 22 were native listeners of Irish English, 2 of American English, and 1 of British English. The study was approved by the School of Psychology Ethics Committee at Trinity College Dublin. All participants signed an informed consent form prior to the experiment. Participants were paid 20 euro for taking part in the study.

### 2.2. Material

#### 2.2.1 Experiment 1 and 2 (L2L and L1L)

Continuous speech sounds were employed in this study (see Data and code availability for stimuli and data). Speech sounds were pre-recorded for this experiment by a female native speaker of British English in the form of storytelling. Each story was recorded in two speech registers: one where the speaker was instructed to address a native English speaker (NDS), and another where she was directed to speak as if addressing a Spanish-speaking novice learner of English (FDS). It’s important to note that the speaker was accustomed to addressing L2 learners (Spanish L1) due to her teaching experience. We measured the vocalic area within the /a/,/i/,/u/ corner vowels (Uther et al., 2007) and speech rate as the number of syllables/second (Hazan et al., 2015; Kühnert & Antolík, 2017). In line with previous literature (Lorge & Katsos, 2019; Piazza et al., under review; 2023; Uther et al., 2007), FDS stories were pronounced with wider vocalic area (∼ 30%) and lower speech rate (∼ 30%) than the NDS stories. The Slow-NDS register was created by applying dynamic time warping to the NDS speech sounds (Müller, 2007), which kept the acoustic features (pitch height, vocalic area, vowel formants) of the NDS stimuli constant but matched speech rate of the NDS stories to the speech rate of the FDS stories (3/2>slope>2/3). This technique aims to find the optimal alignment between two time-dependent sequences, which are warped in a nonlinear fashion to match each other (see Data and code availability to check the audio stimuli).

Duration of FDS and Slow-NDS stories was about 15 minutes each, while the NDS stories had a duration of about 11 minutes, due to higher speech rate. This option was adopted to maintain the same content for the three stories. English multi-talker babble noise (Krishnamurthy & Hansen, 2009) was added to all the stories (+16 dB SNR) to avoid comprehension floor effect for L2L in experiment 1 and ceiling effect for L1L in experiment 2. Babble noise was created in MATLAB 2014b with a custom script by mixing continuous speech streams of 8 British English speakers (Females = 4). A single signal-to-noise ratio that was avoiding such issues was chosen based on an online pilot study. This pilot study tested comprehension of stories from +0 to +20 dB SNR (4 dB steps). The stimuli employed for experiment 1 were also used for experiment 2. It is worth noting that the stories were recorded by a native British English speaker, as this is the pronunciation most commonly taught in Spanish schools (specifically Received Pronunciation; Vilaplana, 2009), making the recordings more understandable to the L2L tested in experiment 1.

### 2.3 Equipment

#### 2.3.1. Experiment 1 (L2L)

Electroencephalography (EEG) data were recorded using a 64 Ag-AgCl electrodes standard setting (two actiCAP 64-channel systems, Brain Products GmbH, Germany) with hardware amplification (BrainAmp DC, Brain Products GmbH, Germany). Signals were bandpass filtered between 0.05 and 500 Hz, digitised using a sampling rate of 1000 Hz, and online referenced to the left earlobe via hardware. PsychoPy 2021 Software (version 2.3; Peirce et al., 2019) was employed to present the stimuli and send synchronization triggers. Triggers were sent to indicate the start of each trial with contingent stimulus presentation and ensure synchronization with EEG recordings.

#### 2.3.2. Experiment 2 (L1L)

Data were acquired from 64 electrode position, digitized at 1024 Hz using an ActiveTwo system (BioSemi B.V., Netherlands). An additional external electrode was placed on participants’ left earlobe for offline referencing. As for experiment 1, PsychoPy Software 2021 (2.3) was employed to present the stimuli and send triggers.

### 2.4. Procedure

#### 2.4.1. Experiment 1 and 2 (L2L and L1L)

All EEG data were collected in dimly lit and sound-proof booths. Stimuli were presented at a sampling rate of 44,100 Hz, monophonically, and at a comfortable volume from Xiaomi Hybrid Mi In-Ear Pro HD headphones. Participants were asked to listen attentively to three stories while EEG signal was recorded. They were asked to sit calmly and upright while looking at a fixation cross, which was presented on the centre of a computer screen right in front of them (at ∼80 cm of distance from their eyes). During the experimental session, participants were presented with one story per speech register, with counterbalanced order across participants. To avoid any effects derived from specific relations between stories and speech registers (e.g., a certain story is more interesting/easier to understand), each story was presented in all the speech registers across participants (with Latin square counterbalanced story-register association). The continuous narration of each story was divided into five consecutive shorter blocks of ∼3 minutes each. At the end of each block, participants were asked 5 comprehension questions (15 questions per story, 45 questions in total). Experimental sessions lasted ∼2 hours including preparation and testing.

### 2.5. Analysis

#### 2.5.1. Behavioural data

Behavioural data were analysed to identify and discard those participants with very low accuracy, who did not pay a sustained level of attention throughout the experiment or who had very low English proficiency (they could not understand most of the stories, N=1). In addition, accuracy based on responses to the questionnaire was used as a proxy of participants’ comprehension. Each question could be scored a finite number ranging between 0 to 1, which respectively represented wrong and correct answers. Most questions required to list multiple answers, which together summed 1 (see Data and code availability for a complete question list). If participants could recall only part of the possible answers (e.g., 1 out 4 elements) for a completely correct answer, they got a fraction score of 1 (e.g., 0.25 points; since 0.25 x 4 = 1). This was done in order to collect finer-grain accuracy score than binomial (correct/incorrect) response.

#### 2.5.2. EEG pre-processing

EEG signal analyses were performed on MATLAB Software (MathWorks, 2021b), using custom scripts, Fieldtrip toolbox functions (Oostenveld et al., 2011), EEGLAB (Delorme & Makeig, 2004), and CNSP resources (Di Liberto et al., 2024). Offline, the data were resampled to 100 Hz and band-pass filtered between 1 and 8 Hz with a Butterworth zero-phase filter (order 2+2). Channels with variance 3 times larger than the channels median variance were rejected. Channels contaminated by noise were recalculated by spline interpolating the surrounding clean channels in EEGLAB. We had planned to discard from the analysis participants with more than 30% of rejected data or more than 4 contaminated electrodes, but no participants were discarded for these reasons.

#### 2.5.3. EEG analysis

The CE of speech in the different registers was estimated by measuring forward models, or temporal response functions (TRF), capturing the linear relationship between continuous stimulus features and the corresponding neural response. TRFs were calculated with the mTRF-Toolbox (Crosse et al., 2016), which implements a linear regression mapping multiple stimulus features to one EEG channel at a time. The regression included an L2 Tikhonov regularization with parameter λ, and was solved through the closed formula β=(*X*^⊤^*X*+λ*I*)^−1^*X*^⊤^*y*, where β indicates the regression weights, *X* the stimulus features, *I* the identity matrix, and *y* an EEG channel. The regularization parameter was selected through an exhaustive search on a logarithmic parameter space from 10^−2^ to 10^8^. This selection was carried out via cross-validation to maximize the EEG prediction correlation averaged across all channels, leading to TRF models that optimally generalize to unseen data. The interaction between stimulus and recorded brain responses is not instantaneous, as a sound stimulus at time *t_0_* can affect the brain signals for a certain time-window [*t_1_*, *t_1_+t*_win_], with *t_1_* ≥0 and *t_win_* >0. The TRF takes this into account by including multiple time-lags between stimulus and neural signal, providing us with model weights that can be interpreted in both space (scalp topographies) and time-lag (speech-EEG latencies).

Stimulus and EEG time-series were split into folds of equal length. Leave-one-out cross-validation procedure was employed to maximize the amount of data used for the model fit, at the cost of additional computational time compared with a single train-test split. Each iteration provided a prediction correlation coefficient (*r-*value) between each feature and the EEG response (per channel). The prediction correlation coefficient is the estimate of how strongly an EEG signal encodes a given set of stimulus features. An *r*-value of 1 would represent perfect correspondence between EEG signal and TRF features, whereas *r*-value of 0 would indicate no correlation whatsoever. It is important to stress that the prediction correlation values (Pearson’s *r*) were extracted from the EEG signal, which is inherently noisy. That is, prediction correlation values have low values that are typically around ∼ 0.05 or ∼0.1 due to the large amount of independent EEG noise and the lack of a ground truth for our evaluation, yet being significant and informative (Brodbeck et al., 2018; Di Liberto et al., 2015, 2021).

#### 2.5.4. Encoding of speech features (TRF regressors)

The CE of speech features of interest was estimated by relating those features with the EEG signal with multivariate TRFs. The stimulus features considered here were the speech envelope, phonetic feature categories, and semantic surprisal. The TRF procedure offers two dependent measures that can be studied to infer the CE of the stimulus. First, the *TRF weights* (i.e., linear regression weights, where a large weight, positive or negative, indicates a stimulus feature and time-lag of particular importance for predicting the EEG signal). Second, *EEG prediction correlations* are derived for each EEG channel, informing us on how informative a feature is for predicting EEG signals at a particular scalp location. Here, TRF models were fit for the three registers (and for each participant) separately, allowing us to compare the CE of speech across different registers with both EEG prediction correlations and TRF weights measures.

##### Speech envelope

The broadband sound envelope was extracted from the speech sounds using the Hilbert transform (Crosse et al., 2021). Univariate TRF models were fit to describe the linear mapping from the speech envelope to the EEG data. We called this the Env model. In this case, the time window used to fit the TRF model was [-200, 600]ms, based on previous research that found this time-lag window to be sufficient to capture the measurable EEG response to the speech envelope (Broderick et al., 2018; Klimovich-Gray et al., 2023).

##### Phonetic features

Phonemic alignments of the speech material were obtained through forced alignment, initially performed automatically using DARLA (Reddy & Stanford, 2015) and subsequently verified manually with PRAAT (Boersma & Weenink, 2001). The alignments were stored as time-series data, with ones marking phoneme onsets and zeros elsewhere. This time-series representation was 19-dimensional, where the different dimensions corresponded to phonetic articulatory features. Phonetic features indicated whether each phoneme was voiced, voiceless (consonants), plosive, fricative, affricate, nasal, liquid, glide, front, back, central, diphthong, close, open (vowels), bilabial, labiodental, dental, alveopalatal, velar-glottal (Ladefoged, 2006). This way, each phoneme could be described as a particular linear combination of phonetic features. A linear transformation matrix was derived to describe the linear mapping from phonetic features to phonemes, which we used to rotate stimulus matrices and TRF weights from phonetic features to the phoneme domain (Di Liberto et al., 2015; Liberto et al., 2021). TRF were fit to describe the mapping of phonetic features to EEG signals. TRF models were fit (with time-lag window [-200, 600]ms) by including phonetic features and acoustic spectrogram (Sgram) simultaneously (PhFSgram multivariate TRF model). Sgram was implemented in 8 frequency bands ranged between 250Hz and 8000Hz. Those bands were defined based on the Greenwood equation that correlates the position of the hair cells in the inner ear to the frequencies that stimulate their corresponding auditory neurons (Greenwood, 1990). Sgram was chosen instead of Env as it provides a richer representation of the speech acoustics, offering in itself acoustic information that can distinguish different phonemes and phonetic features, particularly in terms of frequency variations. For this reason, Sgram expectedly serves a better purpose to control for acoustics than speech envelope when investigating phonetic features. We projected TRF weights from phonetic feature to phoneme domain, and calculated pair-wise Euclidean distances between phonemes for each group and condition. These distances represent how different the encoding of two given phonemes is in the EEG signal (Phoneme distance maps), and it has been associated with language proficiency and nativeness (Di Liberto et al., 2021). These distances can also be visualized by means of a multidimensional scaling analysis (see Supplementary Material figure A. Here, we assessed how the phoneme distances maps are affected by speech register across the two groups.

##### Semantic surprisal

For investigating encoding of semantic surprisal, we first calculated its values as the negative logarithm of the probabilities extracted from the Generative Pre-trained Transformer 2 (GPT-2). GPT-2 calculated the probability of the upcoming words of each sentence of the stories given the previous context. Surprisal values were then coded into a sparse time-vector, where non-zero values represent word onsets, and their values the surprise of that word based on the preceding context. Then, we fit a multivariate TRF including semantic surprisal values and speech envelope as input features (SemEnv TRF model). This was implemented in order to account for the acoustics of speech, while investigating non-acoustic features (Chalehchaleh et al., 2024; Di Liberto et al., 2021). The time window considered for this model was -200 – 700ms based on previous literature showing that EEG responses to semantic surprisal emerges with long latencies (Broderick et al., 2018; Di Liberto et al., 2021; Klimovich-Gray et al., 2023), with longer latencies for L2 than L1 learners (Di Liberto et al., 2021; Momenian et al., 2024; Mueller, 2005).

### 2.6. Statistical analysis

#### 2.6.1. Comprehension questionnaire

To assess the effect of Speech register, statistical analyses were performed using generalized linear mixed effect (*glme*) models with Poisson family, including fixed effect of Speech register, random effect of participants (see Appendix 4.1 for a list of statistical models). To determine significance of the models we used the type II Wald chi-square tests included in the CAR package (Fox, 2015; Fox & Weisberg, 2019). For post-hoc analyses, we used the *emmeans* package with Tukey HSD correction for multiple comparisons.

#### 2.6.2. TRF model performances

Before our analyses of interest, we conducted control tests to assess that the model regressors yielded EEG prediction correlations significantly greater than the null model. Specifically, we assessed whether our features of interest were encoded in the EEG signals to some extent. We conducted one-sample *t*-tests (one-tailed) against the null hypothesis (prediction correlations were not greater than 0) for the speech envelope, phonetic features, and semantic surprisal regressors. Whereas for the Env model we employed the EEG prediction correlation (Pearson’s *r*) of the final model, the phonetic features and semantic surprisal were employed in multivariate models. Thus, we measured the unique contribution of phonetic features and semantic surprisal on model performance by comparing the multivariate models with the EEG prediction correlations for univariate models (Sgram and Env respectively) and subtracted the *r*-values of those from that of the multivariate models. Thus, we assessed whether the residual *r*-values were greater than 0 (Di Liberto et al., 2018; Di Liberto et al., 2015). While there are caveats to this procedure (Daube et al., 2019), this was sufficient for our purposes of studying the impact of speech register across L1L and L2L groups.

#### 2.6.3. TRF regression weights

*Speech envelope.* After assessing model performances, we investigated the effect of Speech Register on the TRF weights of the Env model via the TRF N1-P2 complex (peak-to-peak) amplitude metric. This choice is in line with previous research and recommendations (Carta et al., 2023; Di Liberto et al., 2018, 2021). Thus, we picked the most negative and positive values of each electrode in the 30-180ms TRF time window and calculated the N1-P2 TRF complex of the Env model as the peak-to-peak difference (Crosse et al., 2016; Di Liberto et al., 2015; Di Liberto et al., 2021). We then fitted an linear mixed effect (*lme*) model with Speech Registers as fixed effects and participants as random effects, whereas significance was assessed via type II Wald chi-square test.

##### Phonetic features

We derived the phoneme distance maps as described above by employing multidimensional scaling (MDS) to project the TRF phoneme weights onto a multidimensional space for each speech register. The result for each speech register was then mapped to the average English Listeners’ NDS space by means of a Procrustes analysis (MATLAB function *procrustes*). Then, we calculated residual distance between these three L2L phoneme representations and the reference maps of NDS-L1L (as in Di Liberto et al. 2021). This analysis allowed us to project the L2 phoneme distance maps for the three speech registers’ perception to a common multidimensional space and to compare them quantitatively. The results of the MDS were then fitted to a *lme* model (including participants as random effects) to assess the difference across speech registers in the L2L group.

##### Semantic surprisal

From the SemEnv model, only the temporal weights of the semantic surprisal feature (excluding Env) were analysed at the electrode level employing CBPT. We implemented this approach to avoid selecting a priori time window, as previous literature showed various time windows of the N400 TRF complex (Broderick et al., 2018, 2021; Klimovich-Gray et al., 2023).

The exact same pre-processing and analysis of experiment 1, both for behavioural and EEG data, were conducted on the L1L data of experiment 2. Even though all these steps overlapped between experiment 1 and 2, two separate analyses were conducted because data were collected in two different laboratories and with different EEG recording systems (Brainvision and Biosemi). In Experiment 2, the PhF regressor was primarily used to create L1L- NDS phoneme maps for studying L2 phoneme perception in Experiment 1. We confirmed that the PhF regressor in the PhFSgram model produced a significant EEG prediction correlation (*t* = 9.624, *p* < .001), indicating phonetic feature encoding by L1 listeners, but no further analysis was conducted.

## 3. Results

### 3.1. Speech envelope model

*L2L.* We examined the EEG results of the speech envelope (Env) model performance and whether Env TRF weights differed across speech registers (measured on N1-P2 complex, the ERP equivalent is widely used in the literature; Lightfoot, 2016). Encoding univariate TRF model Env yielded prediction correlations that were higher than zero (*t* = 77.391, *p* < .001), demonstrating that speech envelope was encoded in the EEG signals. The N1-P2 complex yielded a statistically significant effect of speech register (χ^2^ = 408.99, *p* < .001; Figure 1A-C). Post-hoc analyses showed larger N1-P2 complex for FDS than NDS (β = 25.9, *z* = 9.569, *p* > .001) and Slow-NDS (β = 54.8, *z* = 20.214, *p* < .001), which in turn exhibited reduced amplitude as compared to NDS (β = 28.8, *z* = 10.645, *p* < .001).

**Figure 1.**
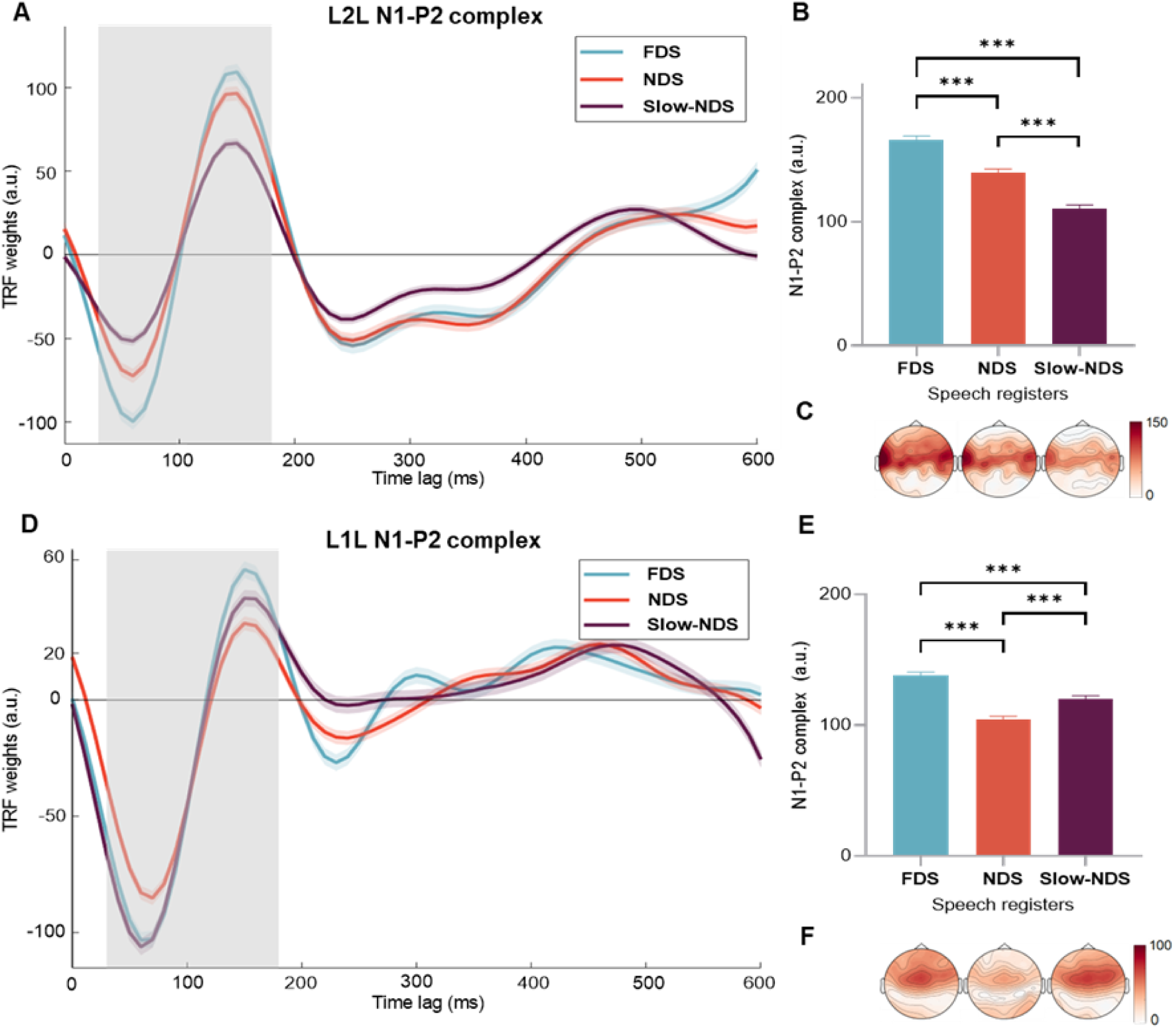
Slow speech rate alone does not support acoustic processing. *A-C: L2L. D-F: L1L.* **(A)** Mean TRF weights of the speech envelope model by Speech Register (FDS = Foreigner- Directed Speech, NDS = Native-Directed Speech, Slow-NDS = Slow-Native-Directed Speech) for Cz channel at post-stimulus time latencies from 0 to 600ms. Shaded lines indicate SEM across participant (on Cz). The grey area indicates the time window where the N1-P2 complex was measured. **(B)** Mean N1-P2 peak differences by Speech registers. Bars indicate SEM, asterisks indicate significant difference (*p <.05, **p < .01, ***p < .001). **(C)** Topographic distribution of the mean differences. **(D-F)** Same as A-C, but results of the L1 listeners.

*L1L.* Then, we repeated the same analysis on the L1L participants. The model performance one-sample *t*-test showed the Env model yielded prediction correlations higher than zero (*t* = 70.952, *p* < .001). The statistical model revealed a statistically significant effect of speech register (χ^2^ = 274.35, p < .001). Post-hoc analyses indicated a larger N1-P2 complex amplitude when listening to FDS than NDS (β = 33.5, *z* = 11.609, *p* < .001) and Slow-NDS (β = 17.9, *z* = 6.209, *p* < .001), with Slow-NDS showing larger a N1-P2 complex amplitude than NDS (β = 15.6, *z* = 5.400, *p* < .001; see Figure 1D-F).

### 3.2. Phoneme distance maps

*L2L.* We investigated the L2 phoneme encoding results to determine whether the model’s performance accurately reflected phoneme processing, as indicated by prediction correlations greater than zero. Additionally, we investigated whether the phoneme distance maps of L2L listening to FDS were closer to native listeners’ perception of NDS than the other two speech registers. The model performance test (Spectrogram Sgram + Phonetic Features PhF multivariate model – Sgram) yielded prediction correlations greater than zero (*t* = 10.751, *p* < .001). We took the L1L EEG signals in NDS as a reference to build the model for the L2L responses to phonemes. The model, fitted on multi-dimensional scaling (MDS) results of phoneme (Ph) TRF weights, highlighted a significant main effect of Speech Register (χ^2^ = 22.18, *p* < .001). Post-hoc analyses indicated that L2L exhibited phoneme representations closer to L1L (NDS) phoneme perception when exposed to FDS as compared to NDS (β = - 5.237, *z* = -4.468, *p* < .001) and Slow-NDS (β = -4.505, *z* = -3.229, *p* = .004) and no difference between the latter two (β = 0.732, *z* = 1.39, *p* = .859; see Figure 2 and Figure A in Supplementary Material for more detailed plots).

**Figure 2.**
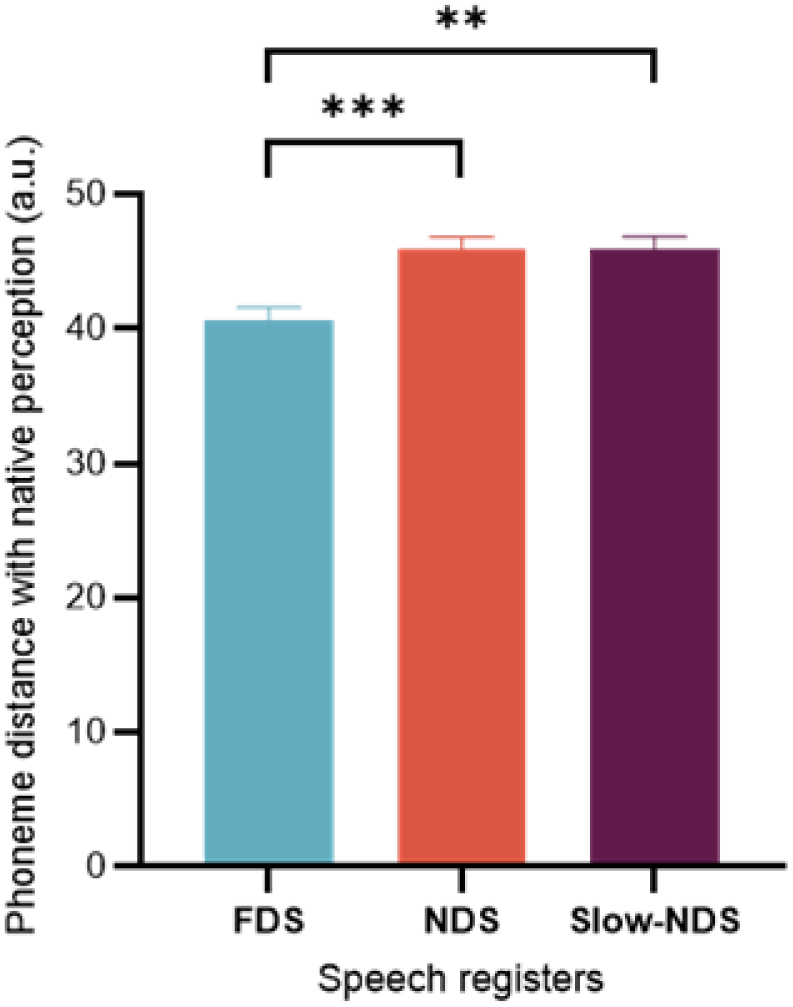
FDS refines L2 phonological encoding. Phoneme distance maps based on the TRF Ph weights. Distance between English listeners’ NDS and L2Ls’ phonemes for each speech register. Error bars indicate the SEM of the mean across phonemes. Asterisks indicate significant differences (*p <.05, **p < .01, ***p < .001).

### 3.3. Comprehension questionnaire

*L2L.* Comprehension scores were compared across the three speech registers via *lme* models. The model revealed a significant effect of Speech register on L2L’s comprehension accuracy (^2^ = 34. 685, *p* < .001). Post-hoc analyses indicated that L2L exhibited higher comprehension scores in FDS than NDS (*z* = 4.793, *p* < .001) and Slow-NDS (*z* = 5.318, *p* < .001), whereas the latter two did not significantly differ (*z* = 0.530, *p* = .857; Figure 3A).

**Figure 3.**
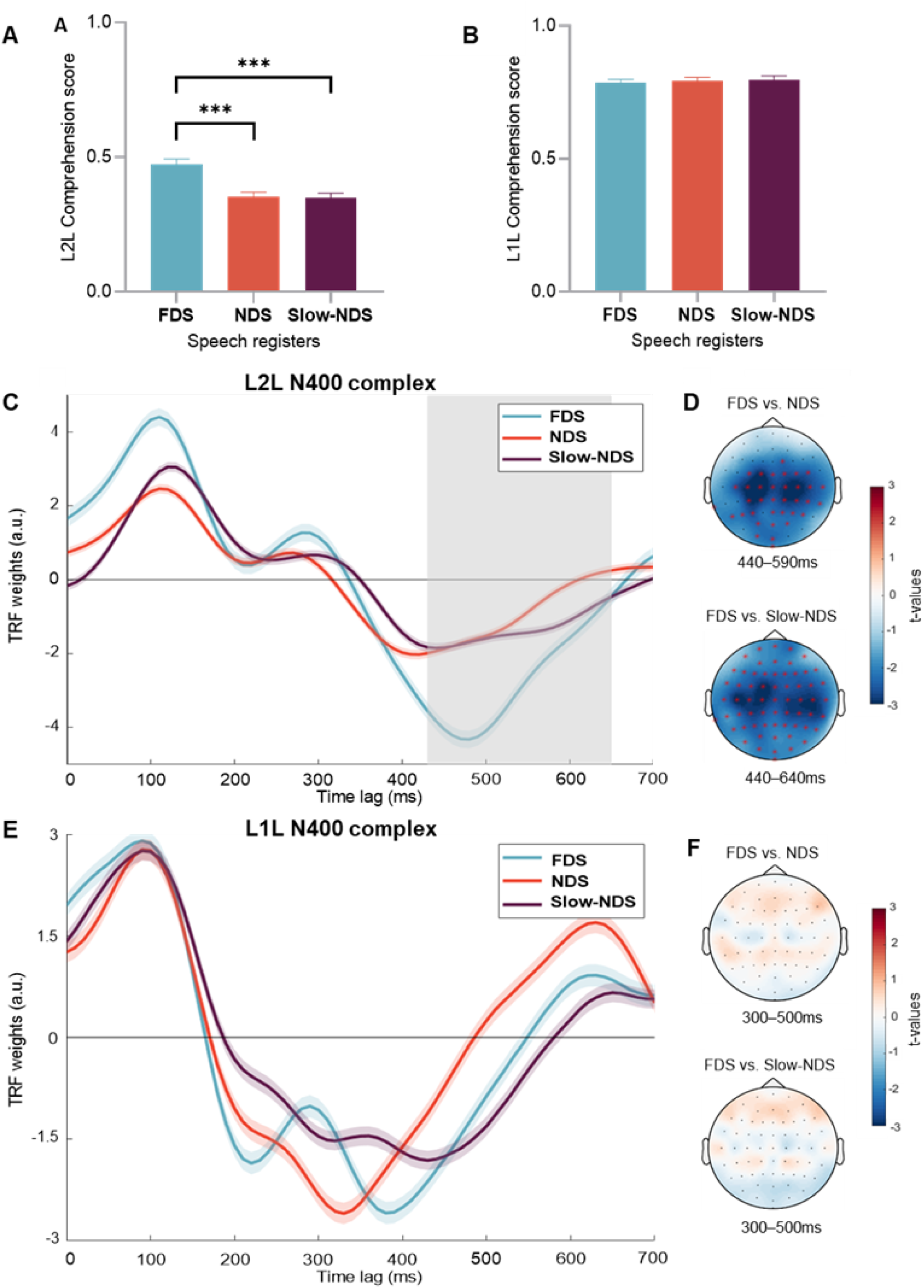
FDS promotes L2 learners’ comprehension accuracy and semantic encoding. **(A) L2Ls** and **(B) L1Ls**’ mean comprehension score by Speech Register (FDS = Foreigner- Directed Speech, NDS = Native-Directed Speech, Slow-NDS = Slow–Native-Directed Speech). Bars indicate SEM. Asterisks indicate significant differences (\**p* <.05, \*\**p* < .01, \*\*\**p* < .001). **(C) L2L’s** mean TRF weight of semantic surprisal (*SemEnv* model) by Speech Registers for Cz channel at post-stimulus time latencies from 0 to 700ms. Shaded lines indicate SEM across participants (on Cz) and grey area indicates the significant time window (CBPT). **(D) L2L’s** significant channels (red asterisks) and time windows of pairwise comparison resulting from the CBPT (difference between speech registers). The NDS vs. Slow-NDS comparison is non-significant and its topography is not reported here. **(E-F)** Same as C-D but results of the **L1L**. The time window to report the topographies is picked arbitrarily as the CBPT did not highlight any significant difference. The NDS vs. Slow-NDS comparison is non-significant and its topography is not reported here.

*L1L.* The statistical model did not yield a significant effect of Speech register (^2^ = 0.495, *p* = 0.976), suggesting that L1L’s comprehension did not benefit from exposure to any of the speech registers (see Figure 3B). L1L response accuracy ranged between 66,7% to 91,6% hinting a lack of ceiling effect.

### 3.4. Semantic surprisal model

*L2L.* Next, we examined whether the difference in register had an effect on the EEG encoding of semantic information. To test this, we fitted a multi-variate encoding TRF model including Semantic Surprisal and Envelope (SemEnv) as its features. The predictive performance of the SemEnv model was significantly higher than that of a univariate model built with the Envelope only (Env), suggesting a robust encoding of semantic information in the EEG responses (*t* = 6.506, *p* < .001). To test the differences across speech registers we employed the cluster-based permutation tests (CBPT; see Method for this rationale) on TRF semantic surprisal weights of the SemEnv model. Results showed that participants exhibited more negative amplitude when listening to FDS stories than NDS (cluster *t* = -2040.5, *p* = .006, SD =.001) and Slow-NDS (cluster *t* = -2081.6, *p* = .002, SD = .0004) in the time window corresponding to the N400 complex (respectively 440-590ms and 440-640ms; see Figure 3C- D). We did not measure significant differences between NDS and Slow-NDS (cluster *p* > .05).

*L1L.* Semantic encoding was tested with the same analysis on L1L participants. Also in this cohort, the SemEnv model yielded a significant prediction gain compared to the univariate Env model (*t* = 3.353, *p* = 0.001), indicating that semantic information was encoded by the L1Ls’ EEG signals. In contrast to L2L listeners, the amplitude of the N400-like response in the semantic surprisal weights did not significantly differ across speech registers for L1L listeners (cluster *p* > .05; see Figure 3E-F)

## 4. Discussion

In this study, we tested the hypothesis that FDS promotes L2 perception and comprehension as compared to NDS and to slow-NDS, characterizing how the choice of speech register impacts speech processing across the cortical processing hierarchy. We also hypothesized that the benefit of FDS would emerge in L2L but not in L1L, the speech register being specifically aimed to address L2L. Our hypotheses were grounded in previous work suggesting that, in comparison with NDS, FDS supports various aspects of L2 acquisition, such as improving L2 perception and comprehension during word learning (Piazza et al., 2022, 2023; Uther et al., 2007). In addition, neurophysiology research on L2 processing had never assessed L2L’s perception in any register but NDS (Piazza et al., 2022; Rothermich et al., 2019, 2023). Conversely, Brodbeck et al. (2024) showed that L2 accented speech can facilitate L2 processing. However, while this research investigates non-standard pronunciation, it does not consider different speech registers, such as FDS. This represents a limitation to the generalization of how L2 is processed in a naturalistic context, where the interlocutors adapt their register to each other (Giles, 2016; Lindblom, 1990; Piazza et al., 2022). Here, we measured EEG responses to NDS, Slow-NDS, and FDS in both L2 learners (L2L) and L1 listeners (L1L). Results showed that EEG signals reflected the encoding of all the speech features considered (speech envelope, phoneme maps, and semantic surprisal), with that encoding being substantially impacted by the speech register.

As we had anticipated, the EEG data supported our hypothesis that FDS promotes the cortical encoding of speech in L2L but not L1L. Our results indicate that FDS promotes speech processing in L2L, and that this effect is not due to the slower speech rate of this register. Instead, L1L only exhibit a sensitivity to speech rate, while the other properties of FDS did not impact the encoding of speech. Our results also indicate that Slow-NDS only alters the cortical encoding of sound acoustics, but not the phonological and semantic encoding, in L1 listeners. In sum, these results indicate that FDS (and not Slow-NDS) promotes L2 encoding at both acoustic (speech envelope) and linguistic levels (phonology and semantics; Figure 1-4).

L2L showed more efficient CE of the speech envelope in FDS than NDS. This advantage of FDS could not be attributed to its acoustic salience due to differences in the speech rate or acoustic onset dynamics, since Slow-NDS and NDS have similar, more gentle rise times compared to FDS (Figure B in Supplementary Material). L1L also showed an effect of register but, contrarily to L2L, they showed significantly enhanced N1-P2 envelope responses due to a slower speech rate, which is in line with previous findings on L1 speech perception (Kösem et al., 2018; Verschueren et al., 2022). One challenge is that the envelope encoding can reflect a variety of contributions and processes, from sound encoding to attention (Ding & Simon, 2012) and even lexical prediction (e.g., attention and engagement; Broderick & Lalor, 2020; Hamouda, 2013). For example, the slower signal in Slow-NDS may be encoded differently acoustically, or it may require less attention, as Slow-NDS is likely easier to process than NDS. And this could be differently reflected on speech envelope CE in L1L and L2L. Note that Slow-NDS was artificially created, but the L1L’s N1-P2 results suggest that it is not perceived as unnatural (Slow-NDS elicited responses comparable to FDS and greater than NDS). Furthermore, no participants reported any issues with its naturalness and audio stimuli are available for verification (Data and code availability section). Importantly, we included Slow-NDS to disentangle the effects of low speech rate on cortical encoding of our regressors. Our results highlight that slow speech rate, if not accompanied by other acoustic features tailored to L2L (as in FDS), is not enough to boost L2Ls’ perception and does not help at any level of the encoding hierarchy.

To more directly uncover the impact of speech register on the encoding of speech, our research probed the cortical encoding of phonological information, providing direct evidence for an enhanced phonological processing among L2L exposed to FDS. Remarkably, our findings revealed that L2L exposed to FDS (as compared to the other 2 registers) exhibited not only better acoustic encoding, but also phoneme perception closer to that of native speakers listening to NDS. In other words, FDS is not merely endowed with more salient acoustics than other speech registers (Hazan et al., 2015; Knoll & Scharrer, 2007); instead, the resemblance of L2Ls’ FDS phoneme maps to L1 NDS maps suggests that L2L exhibit improved phoneme recognition, approaching more native-like perception skills. This represents direct evidence of the FDS impact on phoneme processing in L2L, isolating this factor from purely acoustic factors (spectrogram) and the potential impact of FDS on acoustics. The phoneme map distance analysis allowed us to compare L2Ls’ phonological perception space across three speech registers and contrast it with native perception of NDS. The reason for focusing on NDS perception in L1L as a reference is that it is the register L1 listeners hear daily during peer-to-peer conversations (not FDS), and they have no difficulty to perceive phonemes in this register.

By accounting for both acoustic and phonological features with multivariate TRFs, our analysis could determine that FDS impacts L2 perception and comprehension beyond its acoustic benefit. This approach was previously tested in adults (Di Liberto et al., 2015), children (Di Liberto, et al., 2018b), hearing impaired listeners (Carta et al., under review), L2 listeners (Di Liberto et al. 2021), and infants in their first year of life (Di Liberto et al., 2023), leading to EEG indices of phonological processing that are sensitive to factors such as phonological awareness, language development, proficiency in a second language, comprehension (Di Liberto et al., 2018a), and native vs. non-native encoding of a language. Our finding sheds light on the L2 acquisition process and aligns with existing research supporting the efficacy of FDS in L2 acquisition, including perception of phonemic contrasts (Kangatharan et al., 2023; Piazza et al., 2023; Uther et al., 2012).

On the comprehension-semantic level, we found evidence that FDS promotes L2 comprehension. We observed that L2L had a higher comprehension accuracy to questions about the FDS stories than stories in the other two registers. Additionally, the mTRF analysis was designed to probe semantic prediction mechanisms while accounting for potential contamination from EEG responses to speech acoustics. In the L2L group, we found a modulation of encoding of semantic information in FDS register with more negative N400 TRF complex than in the other conditions (in line with previous research, e.g., Broderick et al., 2018; Klimovich-Gray et al., 2023). This indicates that semantic integration improves when L2L are exposed to FDS, supporting our hypothesis. Notably, the N400 peaked around 500ms, consistent with previous findings and appearing later than typically observed in L1 listeners (Di Liberto et al., 2021; Klimovich-Gray et al., 2023). On the other hand, L1L did not benefit from any register in their comprehension scores. This behavioural null effect was unlikely due to ceiling effects (average accuracy was ∼80%, ranged between 66,7% and 91,6%). Additionally, the semantic surprisal model did not highlight differences across speech registers for L1L, which again suggests no semantic integration advantage in any register. Although L1L acoustic perception was boosted by FDS (speech envelope model), their ability to respond correctly to content questions, and also encode semantics, was not modulated by any register. Also, experiment 2 with L1L employed L1 speakers of other English varieties (mostly Irish, see Method) but the stimuli were presented in British English accent (see Method). We do not think that this negatively affected our results. In fact, L1Ls’ comprehension scores were high (ℒ 80%) and, given the proximity between Dublin and England, contact with accents such as British accent is highly frequent (especially at the University). Thus, we provided evidence that L2L’s – and not L1L’s – comprehension and semantic encoding was boosted by listening to FDS. In our view, such an advantage in L2L’s semantic processing is likely hierarchically linked to the phonological benefits of FDS, in a way that improved phonological encoding facilitates semantic integration. Altogether, these findings support our view that FDS promotes hierarchical speech encoding. Accordingly, speech registers, originated from speech accommodation, promote the intended listeners’ CE of speech at both the acoustic and linguistic levels.

Speakers are known to adapt their speech based on factors like listeners’ language proficiency and communicative intention (Lam et al., 2012; Piazza et al., under review; Rothermich et al., 2023). Theoretical frameworks such as the Communication Accommodation Theory (CAT; Giles, 2016; Giles et al., 1991; Zhang & Giles, 2017) delve into the sociopsychological processes underlying communication, including L1-L2 interaction. CAT, for instance, assumes *convergence* mechanisms, where verbal and nonverbal cues are adjusted to minimize linguistic differences. Our findings provide evidence that speech accommodation indeed impacts intended listeners’ neurocognitive processing mechanisms. We provide compelling evidence that listeners’ CE of speech is enhanced when L2L are exposed to the speech register specifically intended for them. As it seems, speech accommodation affects the intended listeners’ CE and perception of speech. The reason may be that accommodation is particularly relevant for L2Ls’ perception, given their low L2 proficiency. This emphasizes the significance of considering the relationship between speech register and target audience when investigating L1 and L2 processing and building models of speech communication. Auditory L2 speech perception models should integrate various aspects of speech perception including facilitation derived from speech accommodation.

We think that these findings will inform future research on speech interaction. Communication is a dynamic process wherein speakers and listeners cooperate to ensure successful interaction. It is likely that listeners build models of the interlocutors and continuously adjust these models based on contextual information (in line with similar assumptions, Costa et al., 2008; Martin et al., 2016) to maximize communication success. Future research should explore the neurocognitive mechanisms underlying these ongoing adaptation processes and whether this is reflected in CE measures.

## 5. Conclusion

This study on cortical encoding of speech showed that FDS supports L2 learners’ speech processing and comprehension. We highlight the importance of adapting the speech register to the target audience and demonstrate the differential effects of FDS and NDS on language processing in both L2 and L1 listeners. That is, L2 learners process both semantics and phonemes better when they are exposed to FDS than to other speech registers. This study indicates that the speech register employed during communication significantly impacts the degree to which listeners engage and process speech information. These findings have implications for language learning and teaching, and the field of speech communication, emphasizing the significance of tailoring language input to the intended audience.

## 6. Data and code availability

Audio stimuli, processed data (behavioural data, computed EEG metrics), experiment script, statistical formula and analysis code can be found at https://osf.io/ba3p4/?view_only=960986158dd94b92b3b31cca1839b58f. (Anonymized) EEG data and stimuli will be available at https://cnspworkshop.net/index.html in the Continuous-event Neural Data structure (CND format).

## 7. CRediT statement

G.P.: Conceptualisation, Formal analysis, Investigation, Data Curation, Visualisation, Writing - Original Draft, Funding acquisition; S.C.: Formal analysis, Writing: Reviewing and Editing; E.I.: Investigation, Writing: Reviewing and Editing; J.P.N: Resources, Writing: Reviewing and Editing; M.K.: Conceptualisation, Supervision, Writing: Reviewing and Editing. C.D.M.: Conceptualisation, Supervision, Writing: Reviewing and Editing, Funding acquisition; G.D.L.: Conceptualisation, Supervision, Methodology, Writing: Reviewing and Editing, Funding acquisition.

## 10. Supplementary Material

Additional figures can be found at this link: https://osf.io/ba3p4/?view_only=960986158dd94b92b3b31cca1839b58f.

