## Supplementary Material for "Are you talking to me? How the choice of speech register impacts listeners’ hierarchical encoding of speech"

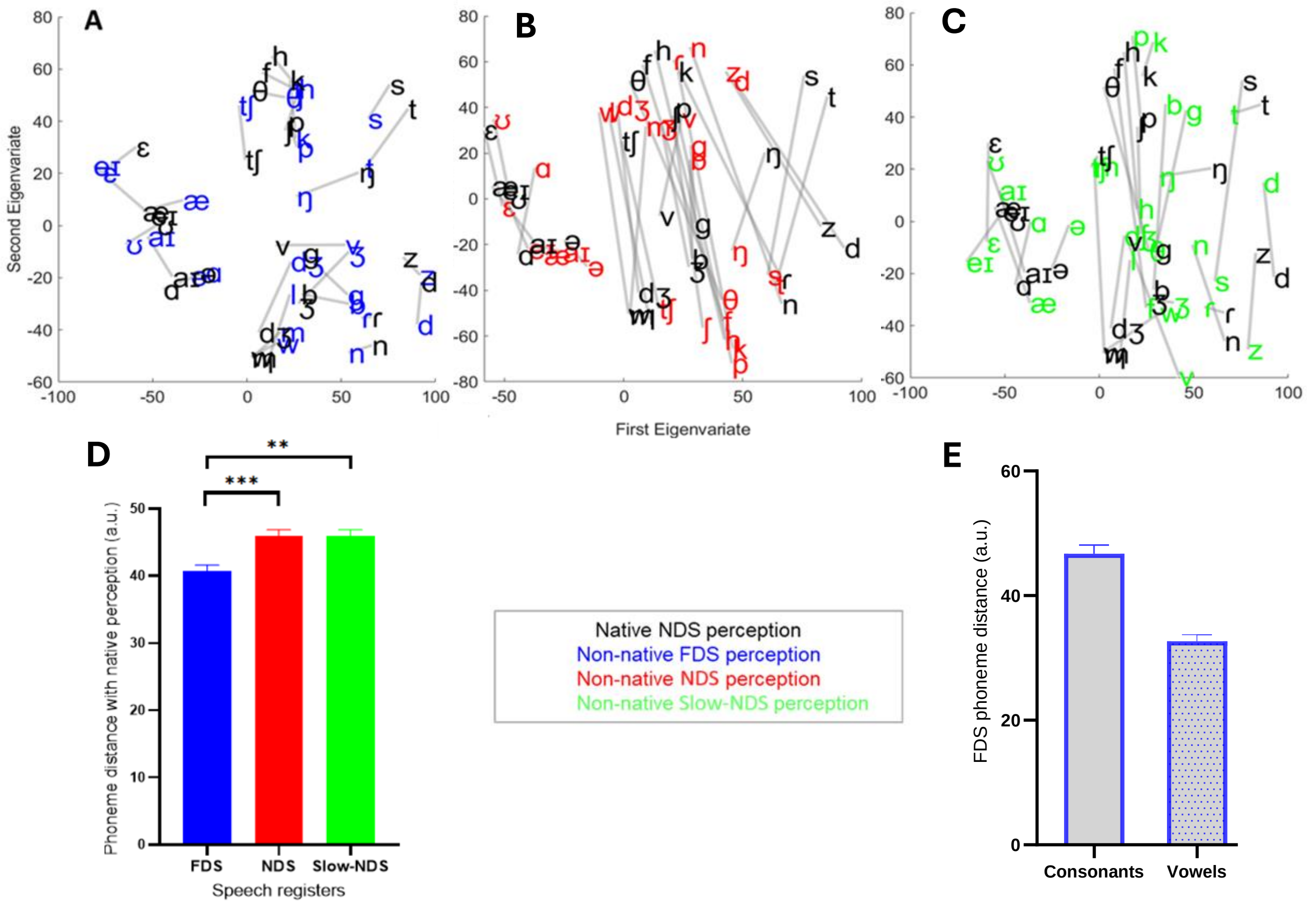


**Figure A. FDS refines L2 phonological encoding.** Phoneme distance maps based on the TRF Phoneme weights. Blue, red, and green colours respectively indicate phonemes for L2 Learners' **(A)** Foreigner Directed Speech, **(B)** Native Directed Speech, **(C)** Slow-Native Directed Speech**,** whereas black colour represent English listener's Native Directed Speech phonemes. **(D)** Distance between English listeners' NDS and L2Ls' phonemes for each speech register. Error bars indicate the SEM of the mean across phonemes. Asterisks indicate significant differences (*$p$ <.05, **$p$ < .01, ***$p$ < .001). **(E)** Descriptive statistics of the L2Ls' FDS phoneme distance by Consonants and Vowels.

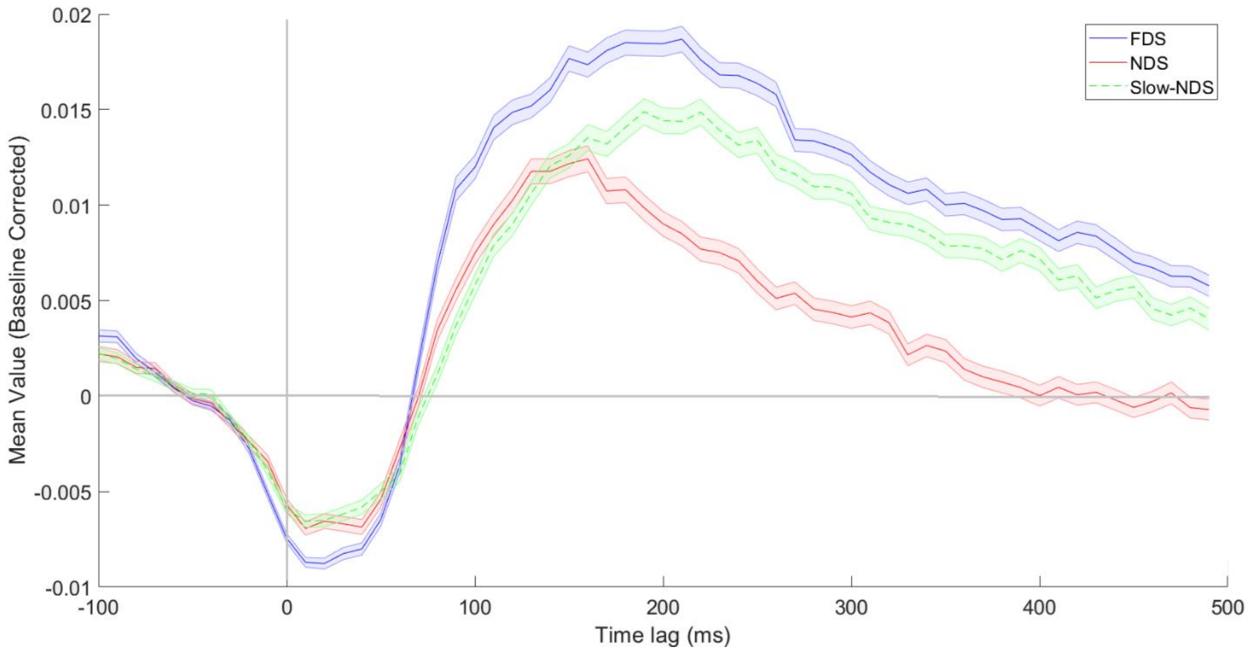


**Figure B.** Mean onset rise time across the three speech registers (Foreigner-Directed Speech, FDS, Native-Directed Speech, NDS, and Slow-Native Directed Speech, Slow-NDS). Shaded area represents SEM.

*Further analysis on the audio-stimuli.*

Amplitude Modulation Spectrum analysis (Goswami et al., 2002, 2010; Pérez-Navarro et al., 2022) revealed that FDS was characterised by local temporal organisation with amplitude modulation bands aligning mostly along the delta frequency band at ~3 Hz (see Figure C). Slow-NDS was pronounced with similar amplitude modulation at ~3 Hz, whereas NDS was mostly aligned ~4.2 Hz. All speech registers rapidly decreased their power (dB below 0) at higher frequency-bands than theta.

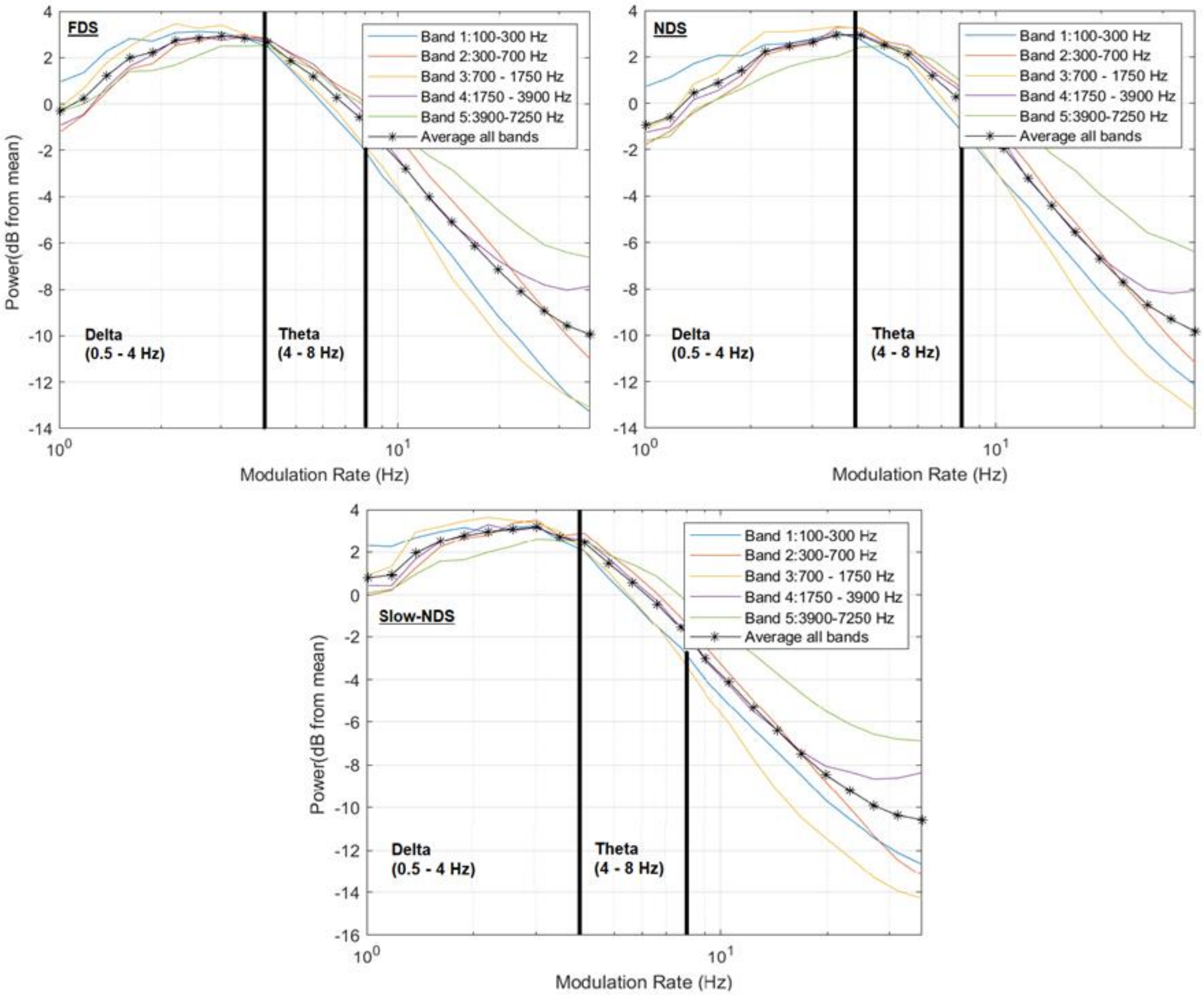


**Figure C.** Amplitude Modulation Spectrum of the experimental stimuli by the three speech registers (FDS = Foreigner Directed Speech, NDS = Native Directed Speech, Slow-NDS = Slow-Native Directed Speech).
